# Parallel Evolution of Key Genomic Features and Cellular Bioenergetics Across the Marine Radiation of a Bacterial Phylum

**DOI:** 10.1101/307454

**Authors:** Eric W. Getz, Saima Sultana Tithi, Liqing Zhang, Frank O. Aylward

## Abstract

Diverse bacterial and archaeal lineages drive biogeochemical cycles in the global ocean, but the evolutionary processes that have shaped their genomic properties and physiological capabilities remain obscure. Here we track the genome evolution of the globally-abundant marine bacterial phylum Marinimicrobia across its diversification into modern marine environments and demonstrate that extant lineages have repeatedly switched between epipelagic and mesopelagic habitats. Moreover, we show that these habitat transitions have been accompanied by repeated and fundamental shifts in genomic organization, cellular bioenergetics, and metabolic modalities. Lineages present in epipelagic niches independently acquired genes necessary for phototrophy and environmental stress mitigation, and their genomes convergently evolved key features associated with genome streamlining. Conversely, lineages residing in mesopelagic waters independently acquired nitrate respiratory machinery and a variety of cytochromes, consistent with the use of alternative terminal electron acceptors in oxygen minimum zones (OMZs). Further, while surface water clades have retained an ancestral Na^+^-pumping respiratory complex, deep water lineages have largely replaced this complex with a canonical H^+^-pumping respiratory complex I, potentially due to the increased efficiency of the latter together with more energy-limiting environments deep in the ocean’s interior. These parallel evolutionary trends across disparate clades suggest that the evolution of key features of genomic organization and cellular bioenergetics in abundant marine lineages may in some ways be predictable and driven largely by environmental conditions and nutrient dynamics.

## Main Text

A fundamental question in microbial ecology concerns the factors that shape the genomic organization and coding potential of bacteria and archaea in the biosphere. With the recent discovery that large swaths of the Tree of Life comprise uncultivated microbial groups that play a central role in shaping the chemical environment of the planet^1,2^, it has become particularly important to expand our knowledge of the evolutionary processes that have given rise to extant microbial life. To this end we analyzed the evolutionary genomics and biogeography of the candidate phylum Marinimicrobia, which has been considered to be a paradigmatic example of “microbial dark matter” because it comprises a diverse group of cryptic microbial lineages that are abundant in the biosphere, and no representative has yet been brought into pure culture and analyzed in the laboratory^3^. The first reports of Marinimicrobia were made in the Sargasso Sea and waters near the Oregon coast, where this group, also referred to as SAR406 or Marine Group A, was identified as a prevalent marine bacterioplankton group distantly related to the Chlorobi and Fibrobacteres^4^. More recent work as elegantly shown that members of this phylum are responsible for a broad diversity of metabolic activities in the ocean and likely play a central role in shaping biogeochemical cycles along thermodynamic gradients^5^. Moreover, other studies have shown that Marinimicrobia are present in a broad array of marine environments including cold seep brine pools^6^, coastal “dead zones”^7^, and OMZs^8,9^, and likely mediate key transformations of nitrogen and sulfur throughout the global ocean^8,10^. In contrast to the broad environmental distributions typical of other bacterial phyla, Marinimicrobia are unusual in that the vast majority of known diversity in this group has been observed in marine environments, thereby providing a unique opportunity to explore the evolutionary processes that have shaped this lineage throughout its radiation into the contemporary ocean.

We compiled a set of 218 publicly-available partial Marinimicrobial genomes that had previously been generated using single-cell or metagenomic approaches^2,3–5,7,11,12^ (see methods). Our phylogenetic analysis of these genomes using concatenated amino acid alignments of marker gene sequences yielded 10 major clades that encompass the majority of known diversity in this phylum (Fig. 1a, Figs. S1, S2). Through comparison of this phylogeny with our genome abundance estimates from Tara Oceans samples^13^ we identified seven clades belonged to a single monophyletic group that are prominent in planktonic marine ecosystems from around the globe (clades 1-7) (Fig. 1c). The remaining three basal-branching clades (clades 8-10) appear to have more restricted biogeographic distributions that include methanogenic bioreactors^11^, deep sea brine pools^6^, and oil reservoirs and fields^14^. The structure base of Clades 1-7, with subsequent lineages diversifying into coastal and pelagic planktonic niches throughout the global ocean. Given that clades 1-7 contained the majority of Marinimicrobial genomes, appeared more prevalent in global ocean waters, and exhibited a more well-defined biogeography, we focused our subsequent analyses on these clades.

**Figure 1.**
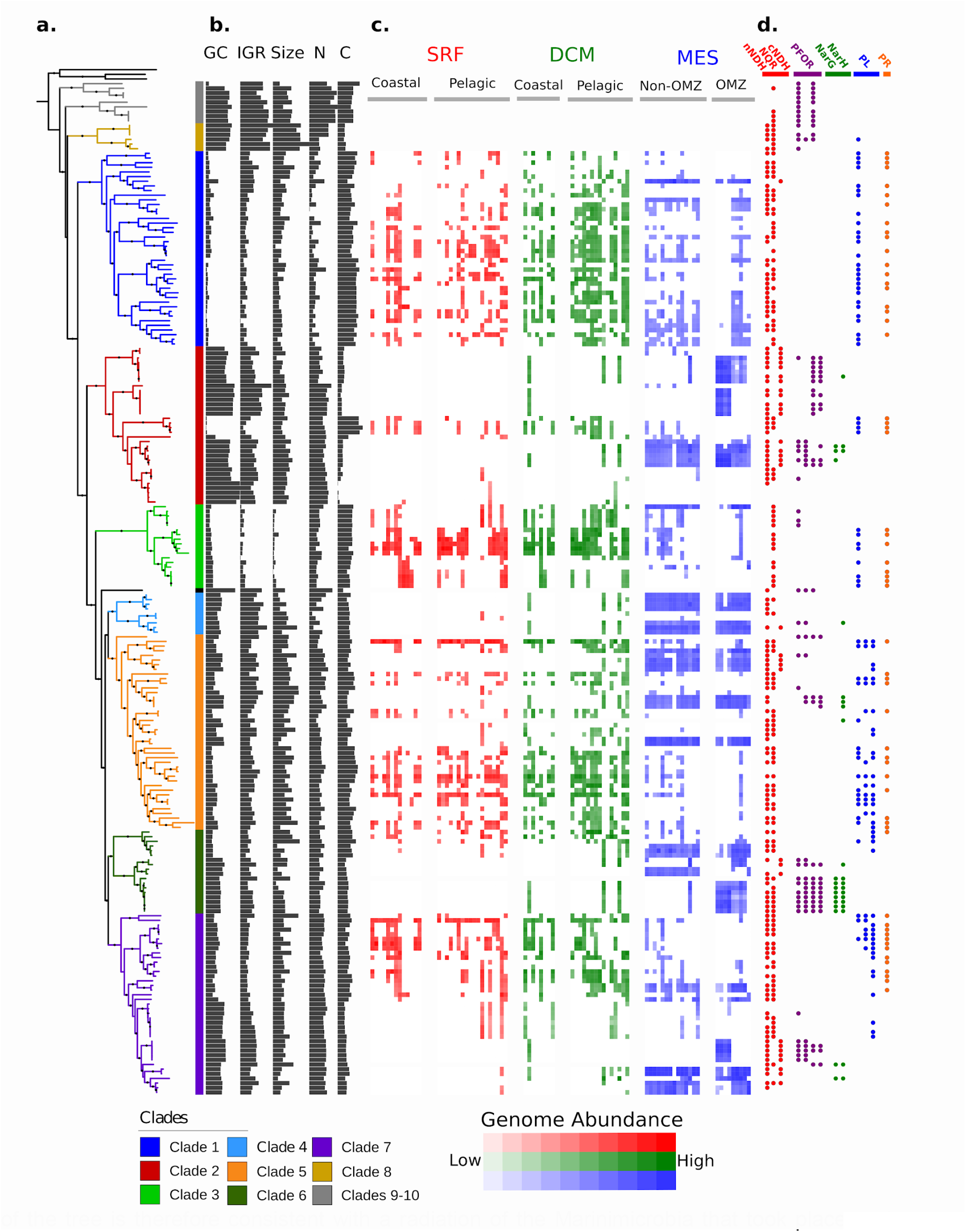
Overview of the phylogeny, genomic features, biogeography, and coding potential of the Marinimicrobia. **A**) A phylogenetic tree of 218 Marinimicrobia genomes constructed using a concatenated alignment of 120 conserved marker genes. Prominent clades are colored, and nodes with support values > 0.95 are denoted with black circles. **B**) Genomic features of the Marinimicrobial genomes. Abbreviations: GC: % GC content (range 30-50%), IGR: Mean intergenic region length (range 40-80), Size: estimated genome size (range 1-3.5 Mbp), N: N-ARSC (range 0.3-0.34), C: C-ARSC (range 3.2-3.4). **C**) Heatmap showing the abundances of Marinimicrobial genomes in different ocean metagenomes. Abbreviations: SRF: Surface Waters, DCM: Deep Chlorophyll Maximum, MES: Mesopelagic, OMZ: Oxygen Minimum Zone. **D**) Presence of select bioenergetic complexes and marker genes in the Marinimicrobial genomes. Abbreviations: nNDH: non-canonical NADH dehyrogenase, NQR: Na^+^-pumping respiratory complex, cNDH: canonical NDH dehydrogenase, PFOR: pyruvate ferredoxin/flavodoxin oxidoreductase, PL: photo-lyase, PR: proteorhodopsin.

We observed that Marinimicrobia in Clades 1-7 were predominantly present in either epipelagic or mesopelagic waters, but not both, consistent with previous findings of distinct structuring of oceanic microbial communities by depth^15–17^ (Fig. 1a,c). We confirmed this finding by clustering genomes of clades 1-7 according to their biogeographic distributions and recovering two major habitat groups that correspond to genomes found in epipelagic or mesopelagic waters (Fig. 2a). While genomes in Clade 3 were found almost entirely in surface waters and genomes in Clade 4 were found almost entirely in mesopelagic waters, genomes in clades 1, 2, 5, 6, and 7 contained several genomes that were found in both environments (Fig 1a,c).

**Figure 2.**
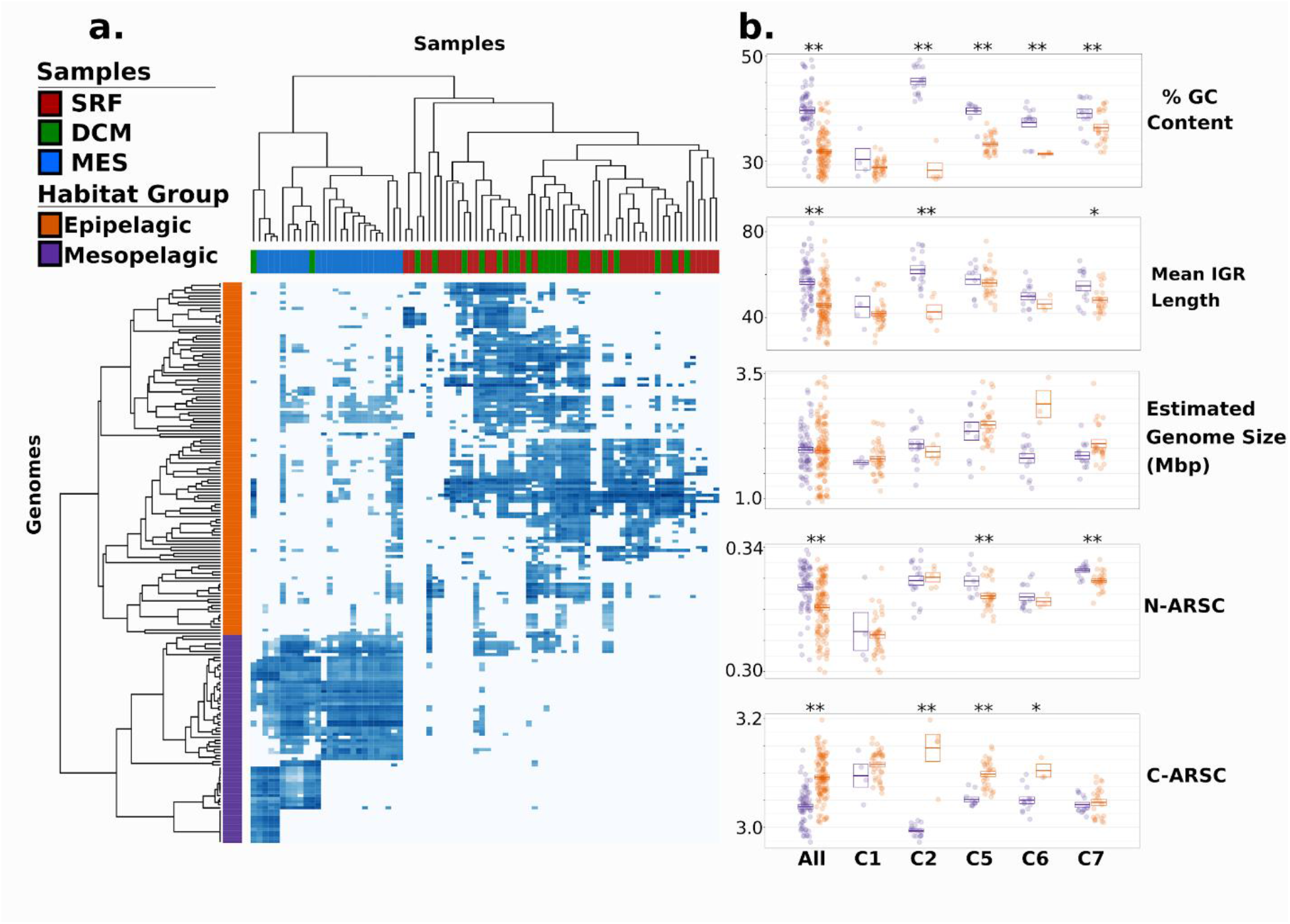
Habitat-based groupings of Marinimicrobial genomes and their genomic features. **A**) Heatmap showing the abundance of Marinimicrobial genomes in Clades 1-7 in different metagenomic samples, with both samples and habitat groups color-coded. **B**) Dot plots showing the genomic features of Marinimicrobia genomes between habitat groups. Each dot is a genome, and boxes provide the means and standard errors. The “All” category includes all genomes in Clades 1-7 that could be assigned to a habitat group, while categories C1, C2, C5, C6, and C7 show only genomes corresponding to those clades. Clades 3 and 4 are not shown since they did not include multiple genomes in both habitat groups.

The biogeographic distributions that we observed throughout the Marinimicrobial tree indicate that several lineages have switched between epipelagic and mesopelagic habitats multiple times throughout their evolutionary history. Concomitant with these habitat switches we observed marked changes in genomic properties, with the genomes of epipelagic Marinimicrobia containing signatures of streamlining such as lower % GC content and shorter intergenic regions^18,19^ (Fig. 1s-c, Fig. 2b). Moreover, epipelagic Marinimicrobia also exhibited fewer nitrogen atoms per residue side-chain in their encoded proteins (N-ARSC), consistent with the hypothesis that this is an adaptation to reduce nitrogen demand in oligotrophic surface waters^15,20^. Conversely, mesopelagic Marinimicrobia contained lower carbon content in their encoded proteins (C-ARSC), consistent with higher nitrogen but lower carbon availabilities in deeper waters^15^ (Fig. 2b). Many of these features were correlated, suggesting the presence of distinct genomic modalities in epipelagic vs mesopelagic Marinimicrobia (Fig. S3), which is consistent with observations of a genomic transition zone (GTZ) between these two regions^15^.

Overall, we found GC content, mean intergenic spacer length, N-ARSC, and C-ARSC to be markedly different between all Marinimicrobia in the two habitat categories (Mann-Whitney U-Test, *p* < 0.01, Fig. 2a), and our intra-clade comparisons demonstrate that these disparities evolved independently in several different clades (Fig. 2b). % GC content was the most prominent feature that shifted with habitat preference, with Clades 2, 5, 6, and 7 all showing significantly lower values in epipelagic vs mesopelagic genomes. C-ARSC was the next most prevalent distinguishing feature between habitat groups, with 3 clades showing significant differences. Interestingly, Clade 1 did not show significantly different genome features between groups despite the epipelagic genomes in this group displaying marked indications of streamlining (Fig. 1). This is likely because only 5 genomes in this clade are more abundant in mesopelagic waters, which limits the statistical power of comparisons. Moreover, the genome with the highest GC content, second highest N-ARSC, and lowest C-ARSC in Clade 1 belongs to Marinimicrobia NORP180, which is the Marinimicrobia in this clade found to be most abundant in the mesopelagic (Fig 1), suggesting that genomic transitions have begun to evolve in this lineage. Other lineages that have only recently switched between habitats may have not had enough time to evolve the genomic features typical of their new environment, indicating that these genomic features require long periods of time to evolve.

Interestingly, we did not observe a consistent reduction in genome size in epipelagic vs mesopelagic Marinimicrobia (Fig. 2b), which is perhaps paradoxical considering the former genomes appeared to more streamlined in all other aspects. Streamlining has been hypothesized to take place over a range of genome sizes, however, since adaptation to a given environment requires particular coding potential that will in turn dictate genome size^18^. Our results affirm this hypothesis and suggest that in the genome streamlining we observe in the Marinimicrobia, which appears to be driven largely by differential nutrient limitations and environmental stressors in epipelagic vs mesopelagic waters, we would not expect to observe a difference in genome size, but rather changes in features such as intergenic spacer length, GC content, and N- and C-ARSC, for which we see strong and repeated shifts.

We also identified clear differences in genomic repertoires between epipelagic and mesopelagic Marinimicrobia, with our pangenomic analyses revealing 758 orthologous groups that were enriched in either of the habitat groups (Fisher’s Exact Test, corrected p-values < 0. 01). Many epipelagic Marinimicrobia in Clades 1-7 have acquired proteorhodopsin proton pumps, photo-lyases associated with UV stress, peroxide stress genes, and phosphate starvation genes, consistent with convergence towards similar mechanisms for life in oligotrophic surface waters in which UV radiation, peroxides, and low nutrient levels are prevalent stressors^21^ (Fig. 3). Our phylogenetic analysis of Marinimicrobial photo-lyases and protoerhodopsins indicate that these genes were acquired independently in different clades, supporting the hypothesis of parallel evolution of distantly related Marinimicrobia towards similar ecological niches (Fig. S4). Conversely, several mesopelagic Marinimicrobia have independently acquired the cellular machinery for nitrate respiration (Fig. 1, S5), mirroring the independent gene acquisitions observed in epipelagic groups and consistent with findings that many Marinimicrobia are poised to exploit alternative electron acceptors under low oxygen concentrations^7,22^.

**Figure 3.**
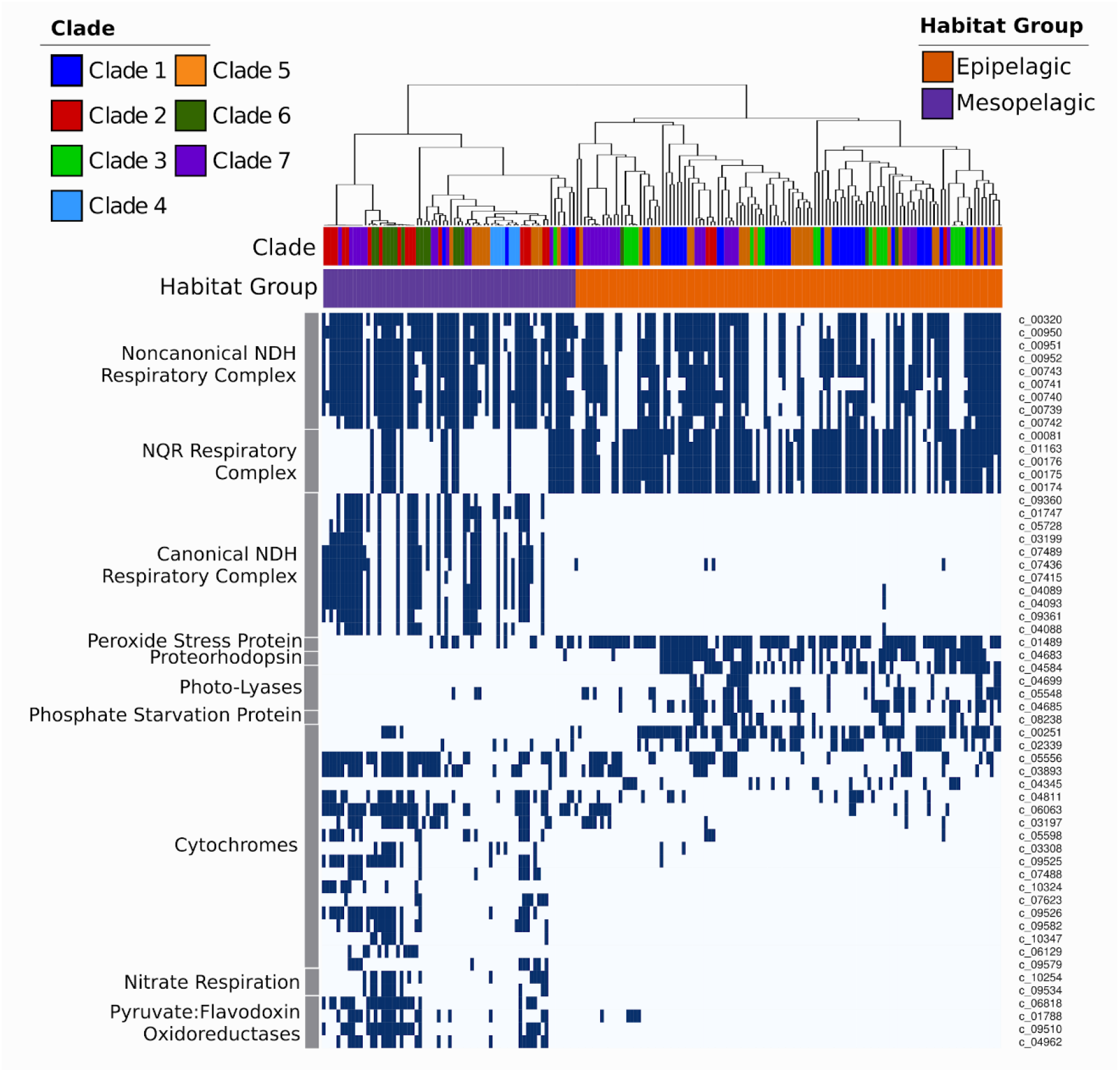
Presence of select marker genes and bioenergetic complexes across the Marinimicrobia. The dendrogram on top shows the habitat-based clustering, and the color strips below it providing the habitat group and clade of the genomes. Unique identifiers for the protein clusters are provided on the right.

We also identified several cytochrome-associated proteins and pyruvate:ferredoxin/flavodoxin oxidoreductases (PFOR) that were differentially enriched in epipelagic vs mesopelagic Marinimicrobia (Fig. 3, Table S1), with all PFOR subunits and most cytochrome subunits more prevalent in mesopelagic groups. Recent work has shown that cytochrome c oxidases are co-expressed with anaerobic respiratory genes in some Marinimicrobia in low dissolved oxygen conditions, suggesting that these cytochromes are either involved in the co-reduction of electron acceptors other than oxygen or involved in the rapid switching between aerobic and anaerobic metabolism^7,15^. Another study of microbial communities in a subseafloor aquifer that included Marinimicrobia also identified genomic signatures of anaerobic respiration despite oxic conditions^22^, further suggesting that switching between electron acceptors may be a dynamic process in deep marine environments that is dictated by prevailing environmental conditions. Overall, the presence of a wide array of cytochromes in mesopelagic Marinimicrobia is consistent with their use of a variety of terminal electron acceptors, similar to what has been observed in other microbes such as *Shewanella oneidensis*^23^. The prevalence of PFORs in mesopelagic Marinimicrobia is also consistent with the metabolic versatility of mesopelagic Marinimicrobia, since the ability to shuttle electrons through alternative carriers such as ferredoxin or flavodoxin may allow for a broader range of respiratory complexes. Our phylogenetic analyses of PFORs are consistent with multiple independent acquisitions by mesopelagic Marinimicrobia (Fig. S6), further indicating convergence towards similar bioenergetic modalities among habitat groups.

Perhaps most strikingly, the different evolutionary forces experienced by epipelagic and mesopelagic Marinimicrobia also appear to have altered their cellular bioenergetics, as we observed a prevalence of NQR-(Na^+^) respiratory complexes in epipelagic Marinimicrobia, while mesopelagic groups appear to have largely replaced this with a canonical NDH-(H^+^) respiratory complex (cNDH) (Fig. 1d, Fig. 3). Most genomes in both groups also encoded a non-canonical NDH-(H^+^) respiratory complex (nNDH) for which the NADH reductase subunits were missing, suggesting that alternative electron donors such as flavodoxin may be used, similar to what has been observed in other groups^24^. These findings indicate that while use of both a sodium motive force and a proton motive force is prevalent across Marinimicrobia, the relative importance of these bioenergetic gradients and how they are used differs between groups. There are a number of reasons that may explain these differences between epipelagic vs mesopelagic Marinimicrobia. Firstly, the canonical NDH-(H^+^) complex is likely more efficient than the NQR- (Na^+^ pump^25^, which is potentially favorable to mesopelagic Marinimicrobia since carbon and energy are less readily available deeper in the water column. For epipelagic Marinimicrobia, an NQR respiratory complex may be sufficient given that nitrogen and phosphorus availabilities and environmental stressors more often limit growth for these bacterioplankton than energy. Secondly, epipelagic waters have slightly higher pH and salinity^26^, which may create a more favorable environment for the harnessing of a sodium motive force in surface waters and a proton motive force deeper in the water column. Lastly, it is possible that the reactive oxygen species (ROS) produced by NDH make the use of this complex disadvantageous in surface waters where high hydrogen peroxide concentrations already generate substantial quantities of these stressors^27^, though it is unclear if the NQR complex of Marinimicrobia produces fewer ROS.

A combination of vertical inheritance and lateral gene transfer (LGT) appears to have shaped the distribution of respiratory complexes throughout the Marinimicrobia. The NQR complex is prevalent throughout the Marinimicrobia phylogeny, including basal-branching Clades 9 and 10, suggesting that this complex was present in the common ancestor of all Marinimicrobia (Fig. 1d). Phylogenetic analysis of NqrA revealed a topology similar to that of the main Marinimicrobia clades, further suggesting that the NQR complex was present in the ancestral Marinimicrobia and has evolved primarily through vertical inheritance (Fig S7). The nNDH complex also appears broadly represented in Marinimicrobia, but its absence in the basal-branching groups 9 and 10 suggests that this complex was either acquired at the last common ancestor of clades 1-8 or it was present in the last common ancestor of all Marinimicrobia and was then subsequently lost in Clades 9 and 10. The evolutionary history of the nNDH complex is broadly consistent with the Marinimicrobial phylogeny, consistent with both of these scenarios. The distribution of the cNDH respiratory complex is the most restricted, with only select mesopelagic Marinimicrobia encoding this gene cluster. Moreover, phylogenetic analysis of NuoH subunit in cNDH revealed a phylogeny inconsistent with the Marinimicrobia phylogeny, suggesting that LGT is largely responsible for shaping the distribution of this gene cluster across the phylum. Among the clade of cNDH NuoH proteins, Clade 2 appears to have the most divergent sequences, suggesting this gene cluster may have been acquired at the common ancestor of clades 2-7 and then transferred between clades afterwards (Fig. S7).

Our combined assessment of the evolutionary genomics and biogeography of the globally-abundant candidate phyla Marinimicrobia has revealed a pattern of parallel genomic, metabolic, and bioenergetic transitions that have occurred in multiple clades concomitant with their shifts between epipelagic or mesopelagic habitats. While epipelagic Marinimicrobia are characterized by streamlined genomes, a NQR Na^+^ respiratory complex, proteorhodopsin-enabled phototrophy, and adaptations to mitigate cellular damage due to UV radiation and other environmental stressors, mesopelagic Marinimicrobia display fewer streamlined features and typically encode a canonical NDH H^+^-pumping respiratory complex, genes for nitrate respiration, PFORs, and various additional cytochrome subunits. In addition to providing key insights into the ecological forces that shape this abundant and globally-distributed bacterioplankton lineage, these genomic, metabolic, and bioenergetic transitions also provide a living record of the evolutionary processes that have given rise to extant Marinimicrobia throughout their diversification into the modern ocean.

Continuing to establish the evolutionary processes that have shaped extant marine microbial groups is critical given that climate change and other more localized anthropogenic disturbances are changing global ocean ecosystems and biogeochemical cycles at an unprecedented rate. For example, both oxygen minimum zones and oligotrophic surface waters in oceanic gyres have been expanding due to climate change^28,29^, thereby fundamentally altering marine biomes and the environments in which microbial life continues to evolve. How these global transitions will in turn shift the ecological and evolutionary trajectories of microbial life is unknown, and establishing the evolutionary drivers that have given rise to the patterns of microbial diversity in the contemporary ocean is a critical first step towards being able to predict the outcome of future changes.

## Materials and Methods

### Compilation of the Marinimicrobia genome set and phylogenetic reconstruction

To complile a preliminary Marinimicrobia dataset we downloaded all genomes from GenBank that were annotated as belonging to the Marinimicrobia phylum according to the NCBI Taxonomy database^30^ on October 15th, 2017. Additionally, we supplemented this datasets with previously-published genomes available in the Integrated Microbial Genomes system (IMG^31^) and two recent studies that generated a large number of metagenome-assembled genomes (MAGs)^12,32^. For the Tully *et al.* study we initially considered all genomes classified as Marinimicrobia as well as all genomes no given a classification. We used checkm to assess the completeness and contamination of the genomes^33^, and continued to analyze only those with contamination < 5% and completeness > 40%.

To confirm that all of the genomes were correctly classified as Marinimicrobia we constructed a preliminary multi-locus phylogenetic tree of all genomes using concatenated alignments of phylogenetic marker genes. To ensure that genomes from phyla closely related to Marinimicrobia were not being erroneously included in this analysis we also included a variety of outgroup genomes from phyla known to be present in the same proximal location as Marinimicrobia in the Tree of Life^2^, which included the phyla Chlorobi, Bacteroidetes, Ignavibacteriae Calditrichaeota, Fibrobacteres, Gemmatimonadetes, Latescibacteria, Zixibacteria, and Cloacimonetes as well as the candidate phyla TA06, UBP1, UBP2, UBP11, WOR-3, and Hyd24-12. For initial phylogenetic assessment we constructed a phylogenetic tree using the checkm bacterial marker set (120 genes^33^), which we refer to here as the checkm_bact set. We predicted proteins using Prodigal v2.6.234 and annotated the protein predictions from each genome through comparison to previously-constructed Hidden Markov Models (HMMs) using HMMER3 with the recommended cutoffs previously reported^33^. The scripts we used for this are publicly-available on GitHub (github.com/faylward/pangenomics/). For alignment and phylogenetic reconstruction we used the ete3 toolkit with the standard_trimmed_fastree workflow^35^, which employs ClustalOmega for alignment^36^, trimal for alignment trimming^37^, and FastTree for phylogenetic inference^38^. The final tree can be viewed in Fig. S1 and via an interactive link on the interactive Tree of Life (iTOL39) (http://itol.embl.de/shared/faylward). Upon analysis of this tree we removed three additional genomes from further analysis because they did not group with other Marinimicrobia (TOBG_SP-359, TOBG_MED-784, and TOBG_RS-789). Additionally, to avoid unnecessary redundancy we removed ten MAGs because they had identical phylogenetic placement with other MAGs generated from the same metagenomic data. In these cases the MAG with the highest estimated completeness was retained. Ultimately we arrived at a final set of 218 Marinimicrobia genomes that we used in subsequent analysis. To construct a final tree for we used the checkm_bact marker gene set and the standard_trimmed_fasttree workflow of ete3, with the genomes of *Fibrobacter succinogenes* S85, *Flavobacterium psychrophulum* FPB101, and *Bacteroides fragilis* YCH46 used as outgroups. This tree can be viewed in Fig. 1 and via interactive link on iTOL (http://itol.embl.de/shared/faylward). We identified major clades of Marinimicrobia through visual inspection of this final tree. Detailed information for all genomes used in this study can be found in Dataset S1.

### Calculation of genomic characteristics

We predicted GC content, N-ARSC, C-ARSC and mean intergenic space length using previously-described methods^15^. Code for these analyses is available online (https://github.com/faylward/pangenomics/). To estimate genome size (S) we used the following formula:

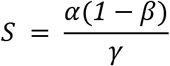

Where *α* is the number of basepairs in the genome assembly, *β* is the estimated contamination and *γ* is the estimated completeness. We estimated contamination and completeness for each genomes using checkM v1.0.7 33.

### Protein cluster identification and annotation

We predicted proteins from all genomes using Prodigal, and subsequently identified protein orthologous groups (OGs) using proteinortho v5.16b with default parameters^40^. For each OG we chose the longest member as a representative compared these proteins to the EggNOG release 4.5^41^, Pfam release 31^42^, and TigrFam release 15.0^43^ databases for annotation using HMMER3^44^. For EggNOG we downloaded all NOG hmms from the EggNOG website on February 1st, 2018, and ran hmmsearch with an e-value cutoff of 1e-5. For Pfam and TigrFam annotations we used the the noise cutoffs in each HMM as lower bounds for annotation.

### Respiratory complex annotation

The canonical NDH (H+) respiratory complex consists of 14 subunits *(nuoA-N)* that correspond to the COG HMMs COG0838, COG0377, COG0852, COG0649, COG1905, COG1894, COG1034, COG1005, COG1143, COG0839, COG0713, COG1009, COG1008, and COG1007. We identified protein OGs that corresponded to these COGs and considered a genome to contain this complex if >= 6 of the the genes from nuoA-L were present, since nuoM and nuoL were found to sometimes be encoded adjacent to other respiratory complexes. We identified a second NDH respiratory complex in many genomes that was lacking subunits D-F, and OGs corresponding to these subunits were distinct from those of the canonical NDH complex. We considered this second NDH complex to be present if >= 5 of the OGs corresponding to the nuoABCGHIJK subunits could be identified. For simplicity we refer to the canonical nuo complex as cNDH and the noncanonical version lacking the nuoDEF as nNDH. The canonical NQR-(Na^+^) respiratory complex consists of the 5 subunits nqrA-F which correspond to COG HMMs COG1726, COG1805, COG2869, COG1347, COG2209, and COG2871. We identified OGs that corresponded to those COGs and considered a genome to encode the NQR complex if >= 3 OGs were present. Detailed information for each protein OG, their annotations, and which genomes encoded members can be found online (figshare.com/projects/Marinimicrobia Pangenomics/30881).

### Marker gene phylogenies

To assess the evolutionary histories of key marker genes in the Marinimicrobia we constructed phylogenies with these genes together with available reference sequences. For each marker gene we identified the NOG to which the OG had been annotated using our EggNOG annotations and then downloaded all proteins belonging to the appropriate NOG on the EggNOG website^41^. The one exception to this was for the proteorhodopsin phylogeny, for which we used the reference sequences available on the MicRhoDE database45. Because reference protein datasets were quite large, we reduced their size by clustering similar proteins using CD-HIT^46^ (default parameters). These reference proteins were then combined with the Marinimicrobia proteins into a single FASTA file, and phylogenies were constructed using the ete3 toolkit^35^, with the standard_trimmed_fasttree workflow. We refer to these as the “full phylogenies” since they included a large number of reference sequences. For ease of visualization we manually selected a subset of reference sequences together with all Marinimicrobia sequences from the full phylogenies and then constructed smaller “subset phylogenies”. Subset phylogenies, with Marinimicrobia proteins colored by clade, are provided in Figs. S4-S7. Full phylogenies are available as interactive trees at http://itol.embl.de/shared/faylward.

### Marinimicrobia genome distributions and habitat distinctions

To identify the biogeographic distributions of different Marinimicrobia genomes in global ocean samples we downloaded 90 metagenome samples from the Tara Oceans expedition that represented a broad array of environmental features, depths, and Longhurstian provinces and mapped reads against the final set of 218 Marinimicrobia genomes. We chose Tara Oceans samples to represent as broad a sampling of environments as possible and include different depths (surface, deep chlorophyll maximum, and mesopelagic), ocean basins, and Longhurstian provinces. Details for the samples chosen are available online (<u>figshare.com/proiects/Marinimicrobia Pangenomics/30881</u>). We mapped reads using FastViromeExplorer^47^, which, although initially intended for identification of viral sequences, includes a rapid and versatile read mapping utility which contains built-in filters to remove spuriously-identified sequences. We report final genome quantifications using the TPM metric48, which corrects for sample size and reference genome length.

For genome clustering we loaded a log_10_-transformed genome TPM abundance matrix into R and calculated pairwise Pearson correlation coefficients for the genomes using the “cor” function. We converted these correlations into distances by subtracting from one, and then clustered the genomes using the “hclust” command in R using average linkage clustering. We clustered samples using the same method. Heatmaps and clustering dendrograms were visualized using the heatmap.2 function in the gplots package.

### Supplementary information and data availability

Phylogenetic trees constructed here are publicly available as interactive trees at http://itol.embl.de/shared/faylward. Supplementary data and all key data products generated as part of this study are publicly available online: figshare.com/projects/Marinimicrobia_Pangenomics/30881

## Acknowledgements

The authors thank Jessica Bryant, Roderick Jensen, Stephen Melville, and Melanie Spero for helpful discussions and code sharing. We acknowledge use of the Virginia Tech Advanced Research Computing center for bioinformatic analyses performed in this study. This work was supported by a Sloan Research Fellowship to FOA.

## Author contributions

EWG and FOA performed analyses. SST, and LZ contributed software and bioinformatic expertise. EWG and FOA wrote the paper.

